# IGF2BP3 remodels the microRNA targeting landscape in MLL-AF4 leukemia

**DOI:** 10.1101/2025.10.10.681741

**Authors:** Lyna E S Kabbani, Shruti Kapoor, Gunjan Sharma, Martin Gutierrez, Zachary T Neeb, Amit Kumar Jaiswal, Alexander J Ritter, Sol Katzman, Jordyn Feldman, Dinesh S Rao, Jeremy R Sanford

## Abstract

Insulin-like growth factor 2 mRNA binding protein 3 (IGF2BP3/I3) is a multi-domain RNA-binding protein required for MLL-AF4–driven leukemogenesis, but its mechanism of action remains enigmatic. We hypothesized from our previous work that I3 amplifies oncogenic gene expression by modulating RNA induced silencing complex (RISC) mRNA interactions. To test this, we performed miR-eCLIP of AGO2, the catalytic RISC subunit, in I3 knock-out (I3KO) as well as control B-cell acute lymphoblastic cell lines (B-ALL) and identified I3-dependent AGO2 binding sites on 111 3′UTRs. Analyzing chimeric miRNA-mRNA reads, we observed differential miRNA occupancy in the I3KO compared to control, including increased targeting by miR-181a, a regulator of leukocyte differentiation. Notably, miR-181a overexpression phenocopied I3 loss, implicating I3 in restricting miR-181a-mediated repression. Biochemical assays confirmed direct competition between I3 and AGO2-miRNA complexes for 3′-UTR binding. Taken together, our results provide a model for how I3 promotes leukemogenesis by antagonizing RISC-mediated repression of oncogenic mRNAs.

## Introduction

B-cell acute lymphoblastic leukemia (B-ALL) disproportionately affects children and adolescents. It is the most frequently diagnosed pediatric ALL subtype and affects around 3,000 children every year in the United States, with the highest incidence being in children aged 1-4 years ^1,2^. B-ALL is characterized by an accumulation of precursor white blood cells, crowding out healthy red blood cells, white blood cells, and platelets, impeding normal hematological and immune function. Rapid and aggressive progression is a hallmark for this ALL subtype. Even though the survival rate for B-ALL has steadily risen since 1975, several challenges remain for effective treatment: Around 90% of childhood survivors will experience adverse health effects due to the aggressive nature of treatments such as chemotherapy and radiation therapy ^3–5^.

Insulin-like growth factor 2 mRNA-binding protein 3 (IGF2BP3) is an oncofetal RBP, strongly expressed during embryogenesis, but undetectable in most mature tissues, with the exception of reproductive tissue such as the placenta (during pregnancy) and testes^6–8^. Aberrant expression of IGF2BP3 in mature tissue is recognized as a biomarker for many different types of cancers, and its expression level correlates with disease severity.^9^ In recent years, our group and others discovered a direct role of IGF2BP3 in hematological cancers^10–13^. IGF2BP3 skews white blood cell lineages towards immature hematopoietic stem and progenitor cells and is required for the function of leukemic stem and progenitor cells *in vivo*^11,13^. Deletion of IGF2BP3 greatly attenuates disease in MLL driven acute leukemia in mouse models, with knock-out mice showing highly reduced mortality and tumorigenesis^11^. Additionally, IGF2BP3 depletion enhances the anti-leukemic effects of the small molecule drug MI-503, such as promotion of cell differentiation, apoptosis and inhibition of cell growth^10^.

For most mRNA transcripts, the 3’ untranslated regions (3’-UTRs) harbor a myriad of *cis*-regulatory elements for post-transcriptional control of gene expression. IGF2BP3 preferentially interacts with 3’-UTRs and promotes RNA translation, stability and degradation, depending on context ^8,14,15^. IGF2BP3 binding sites are enriched near microRNA (miRNA) target sites, suggesting that modulation of the RNA Induced Silencing Complex (RISC)-targeting as a potential mechanism ^13,16–18^. Despite mounting evidence for IGF2BP3 as a driver of leukemogenesis, little is known about the exact molecular mechanisms of how it remodels the gene regulatory landscape in leukemic cells. To test the hypothesis that IGF2BP3 remodels RISC-3’-UTR interactions on a global scale, we mapped *in situ* AGO2 binding sites in the presence and absence of IGF2BP3 using miR-eCLIP. This top-down approach revealed a direct role for IGF2BP3 in RISC target selection.

Taken together, our results suggest that competition between tumor suppressive miRNAs and IGF2BP3, most notably miR-181a and let-7a, controls a proliferative post-transcriptional gene expression program which amplifies leukemogenesis.

## Results

### IGF2BP3 modulates gene expression at multiple levels in MLL-AF4 B-lymphoblastic leukemia cells

Previous studies from our group and others have shown that IGF2BP3 (I3) regulates gene expression at various levels^17–20^. To investigate the impact of I3 in MLL-AF4 translocated B-ALL cells, we established a CRISPR-edited SEM cell line harboring a deletion of I3 (I3KO) and performed enhanced crosslinking immunoprecipitation (eCLIP) of I3 as well as RNA-sequencing (RNA-seq) in control and I3KO cells (Fig.1A, Supp.Fig1A). We performed peak calling analysis for I3 eCLIP in the control using Skipper ^21^, which identified 41,772 I3 peaks, followed by annotation of I3 peaks across genomic features. Although the majority of peaks were found in the CDS (Fig.1A), we observe the highest I3 binding intensity in the 3’-UTR (Fig.1B). To query the impact of I3 on global gene expression, we investigated steady-state mRNA levels in control and I3KO cells (Fig.1C). We discovered a total of 2,227 differentially expressed mRNAs (circles), out of which 256 (11.49%) are I3 targets (purple triangles) according to our eCLIP data. With respect to the I3KO, we found 1,040 differentially upregulated mRNAs out of which 99 are I3 targets, while out of 1,127 downregulated mRNAs, 154 are I3 targets. To assess whether the number of I3 targets among the differentially expressed mRNAs represents a significantly enriched fraction, we performed a Fisher’s exact test and found that I3 mRNA targets are significantly overrepresented (p-value = <0.0001). This result suggested that I3 may have a role in transcriptional control and/or regulates the stability of its targets. Because I3 has known functions in translational control of its RNA targets^18,20,22^, we also investigated steady-state protein levels in control and I3KO using Liquid Chromatography-Tandem Mass Spectrometry (LC-MS/MS) (Fig.1D). We identified 741 differentially expressed proteins (circles) out of which 391 (53%) are bound by I3 at their mRNA transcript according to our eCLIP data (purple triangles). With respect to the I3KO we found 427 downregulated proteins out of which 216 are I3 targets and 314 upregulated proteins out of which 175 are I3 targets. Analogous to our RNA-seq data, I3 targets represent a significantly enriched fraction among the differentially expressed pool of proteins (Fisher’s exact test; p-value <0.0001). This indicates a regulatory role for I3 in the translation of its RNA targets.Together, this data suggests that I3 regulates gene expression at both the RNA and protein level in MLL-translocated B-ALL cells.

**Figure 1.**
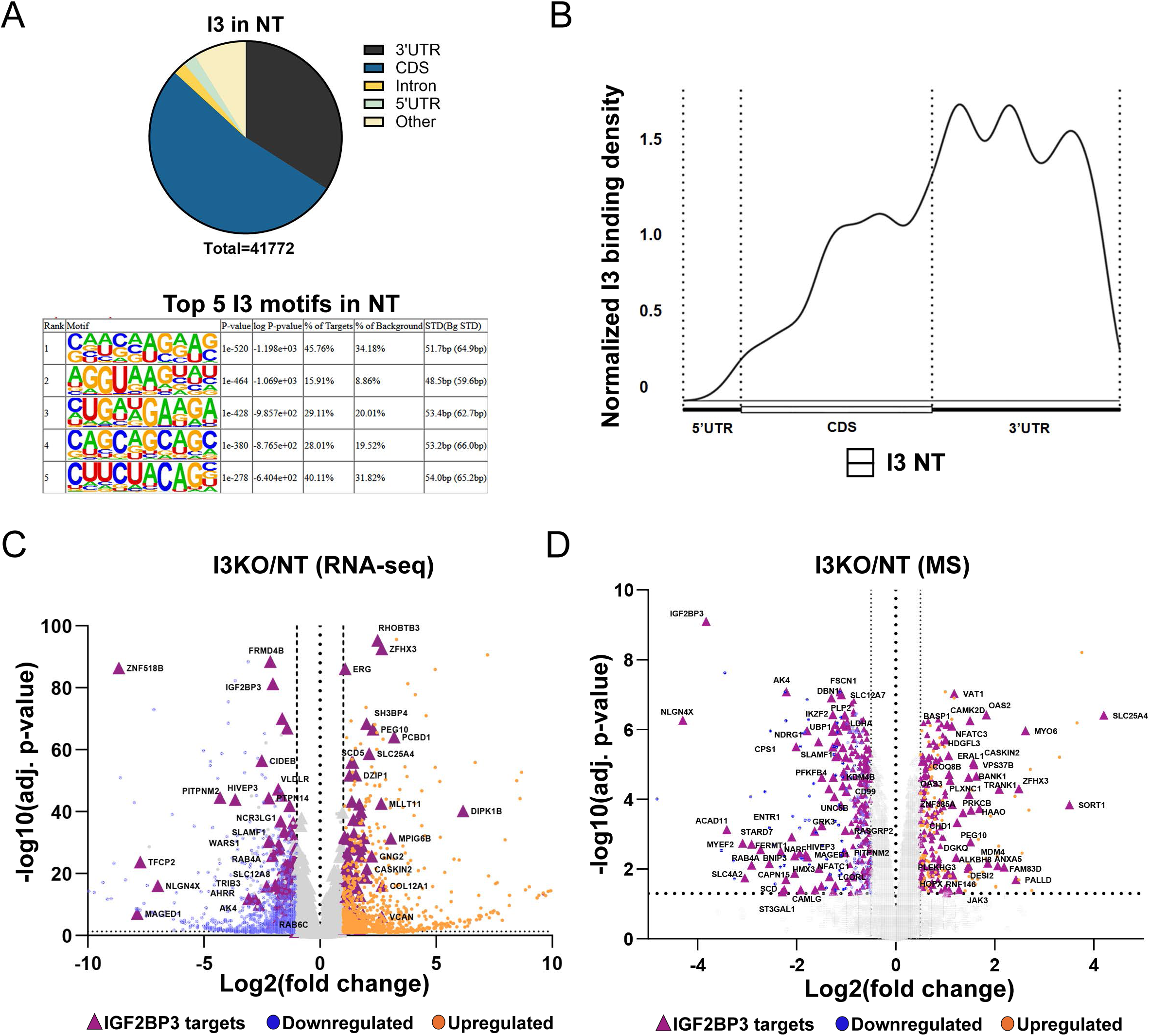
Identification of IGF2BP3 binding sites and differentially expressed genes. (A) Distribution of I3 eCLIP peaks across gene features in control (NT) SEM cells and the top five HOMER binding motifs of I3 eCLIP peaks in control (NT) SEM cells. The total number of eCLIP peaks are listed under the pie chart. (B) Metagene plot depicting the relative density of I3 eCLIP peaks in NT. Peak counts were normalized as counts per million relative to gene length. (C) Volcano plot showing differentially expressed transcripts by RNA-seq between control (NT) and I3 knock-out (I3KO) cells, with respect to I3KO. Significantly differentially expressed RNAs are depicted as colored dots. Transcripts bound by I3 are marked as purple triangles. The threshold for differential peaks was defined as less than -1 Log2 fold-change or greater than 1 Log2 fold-change, with the p-value <0.05. (D) Volcano plot showing differentially expressed transcripts by Mass-spectrometry between control (NT) and I3 knock-out (I3KO) cells, with respect to I3KO. Significantly differentially expressed proteins are depicted as colored dots. Transcripts bound by I3 are marked as purple triangles. The threshold for differential peaks was defined as lesser than -0.5 Log2 fold-change or greater than 0.5 Log2 fold-change, with the p-value <0.05.

### IGF2BP3 alters the AGO2 binding profile on shared mRNA targets and reduces AGO2 binding to 3’-UTRs of genes regulating pro-proliferative pathways

Our results show preferential occupancy of I3 at 3’-UTR sequences and differential gene expression at the RNA and protein level upon I3 knock-out. Data from our previous study in pancreatic ductal adenocarcinoma cells suggests that I3 regulates RISC-mRNA interactions and that I3 is a bi-modal regulator of target mRNA expression^17^.To test this hypothesis we analyzed changes in AGO2 binding in I3KO and control cells. After confirming that loss of I3 does not impact AGO2 protein levels (Supp.Fig.1B) we performed AGO2 eCLIP in control and I3KO cells (Supp.Fig.1C, Supp.Fig.2A-F). After eCLIP peak identification with Skipper, we quantified the number of AGO2 peaks in the I3KO and the control cells (Supp.Fig.2A). While the overall distribution of peaks per gene feature was similar for both conditions (Supp.Fig.2B), we observed an I3-dependent change in the distribution of AGO2 peaks across I3-bound mRNA transcripts (Supp.Fig.2D). Additionally, using HOMER ^23^ to analyze enriched RNA motifs of AGO2, we observed an enrichment for different sets of motif sequences in NT compared to I3KO (Supp.Fig.2C).

**Figure 2.**
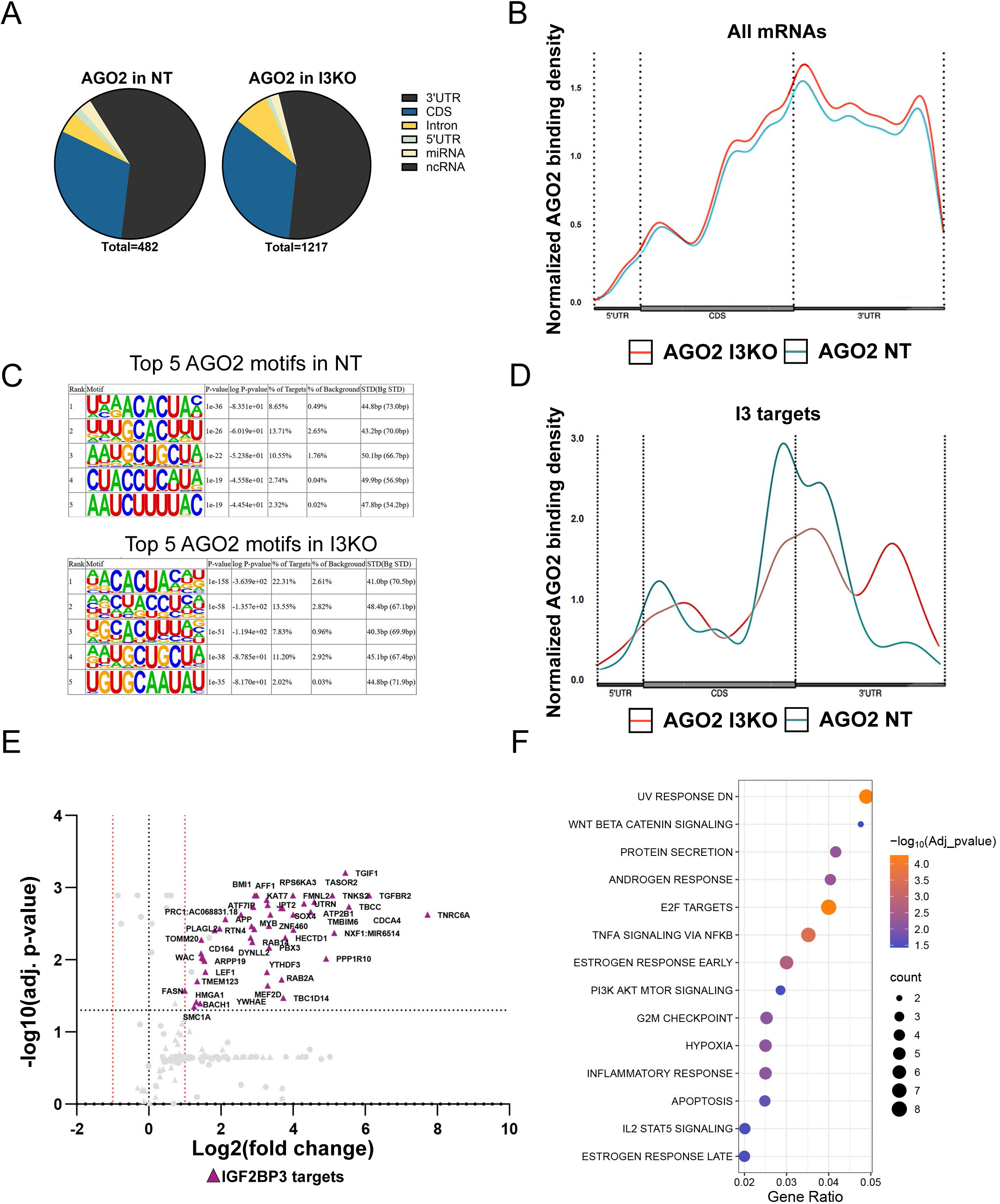
AGO2 binding on oncogenic 3’-UTRs is attenuated by I3. (A) Distribution of AGO2 eCLIP peaks across gene features in control (NT) and I3 knock-out (I3KO) SEM cells. Total number of peaks are listed for each condition under the respective pie charts. (B) Metagene plot depicting the relative density of AGO2 eCLIP peaks across gene features of all mRNAs in control (NT, blue) and I3 knock-out (I3KO, red) SEM cells. Peak counts were normalized as counts per million relative to gene length. (C) Top five HOMER binding motifs of AGO2 eCLIP peaks in control (NT) and I3 knock-out SEM cells (I3KO). (D) Metagene plot depicting the relative density of AGO2 eCLIP peaks across gene features of I3 target mRNAs in control (NT, blue) and I3 knock-out (I3KO, red) SEM cells. Peak counts were normalized as counts per million relative to gene length. (E) Volcano plot showing differential AGO2 eCLIP peaks between control (NT) and I3 knock-out (I3KO) cells, with respect to I3KO. AGO2 eCLIP peaks in 3’-UTRs also bound by I3 in NT are marked as purple triangles. The threshold for differential peaks was defined as less than -1 Log2 fold-change or greater than 1 Log2 fold-change, with the p-value <0.05. (F) Hallmark enrichment analysis of mRNAs with increased 3’-UTR AGO2 eCLIP peaks in IGF2BP3 knock-out cells (purple triangles from panel E). Significantly enriched pathways had an FDR of less than 0.05.

To increase the stringency of our analysis, we subsampled AGO2 eCLIP reads to peaks containing miRNA-mRNA “chimeric” fusion reads^24,25^ (Fig.2). Chimeric reads represent direct miRNA-mRNA connections, while non-chimeric AGO2 eCLIP reads map overall AGO2-RNA associations. We overlapped chimeric peaks with total AGO2 peaks and identified 482 and 1217 AGO2 peaks with chimeric reads in both NT and I3KO samples respectively, with an enrichment towards the 3’-UTR (Fig.2A, Supp.Fig.2H). Analogous to our unfiltered AGO2 eCLIP peaks (Supp.Fig.2B, Supp.Fig.2E) the distribution of peaks per gene feature for all targets was similar for both conditions (Fig.2B); however, for I3 targets, we observe a shift in AGO2 binding density towards the 3’-UTR of I3 targeted transcripts in the knock-out cells (Fig.2D, Supp.Fig.2D). HOMER motifs differ between both conditions (Fig.2C) albeit to a lesser extent than for our unfiltered eCLIP data (Supp.Fig.2C).

To determine if there is a direct relationship between I3 and AGO2-RNA interactions we quantified changes in AGO2 occupancy on 3’-UTRs that were also crosslinked to I3 in control cells (Fig.2E). The volcano plot in figure 2E shows differential peak changes in AGO2 eCLIP on I3-target 3’-UTRs (purple triangles) and non-I3 targets (circles). The majority of I3 target transcripts exhibit a significant increase in AGO2 binding in I3 knock-out cells (Fig 2E). Notably, 47 out of the 111 I3-bound 3’-UTRs show increased AGO2 binding in the I3KO, while none show significantly decreased AGO2 binding. We observe a similar result for our unfiltered AGO2 eCLIP dataset (Supp.Fig.2E). These data suggest that AGO2 binding increases in the I3KO on 3’-UTRs otherwise also bound by I3 in the NT. We call peaks with this pattern “I3-dependent AGO2 binding sites“. Taken together, the I3-dependent AGO2-mRNA interactions suggest the intriguing hypothesis that I3 antagonizes AGO2 binding on 3’-UTRs. Therefore, we wanted to determine if the genes regulated by those 3’-UTRs are relevant to oncogenesis. Following enrichment analysis, we found that mRNAs with I3-dependent AGO2 binding on 3’-UTRs have an enrichment for pro-proliferative pathways (Fig. 2F). Using the Hallmark gene set from the Molecular Signatures Database (MSigDB) we identified several enriched pathways and processes that are linked to cancer hallmarks such as genes regulating apoptosis, cell cycle progression (G2M Checkpoint, E2F Targets), and signaling pathways (TNFA signaling via NF-KB, WNT Beta Catenin, PI3/Akt/mTOR signaling, IL2 STAT5), among others (Fig.2F). Similarly, we also observe enriched cancer-related pathways for 3’-UTRs with I3-dependent AGO2 binding in our unfiltered AGO2 data set (Supp.Fig.2F). Extended supplemental figure 2C shows browser snapshots of two example mRNAs, *HOXA7* and *BCL2L11*. Differential AGO2 peaks between each condition are highlighted. To determine if the increase in AGO2-3’-UTR interactions, which we observed in I3KO cells, occurs at functional sites within 3’-UTRs, we quantified AGO2 peak density relative to annotated miRNA target sites in control and I3KO cells. Interestingly, deletion of I3 resulted in a significant increase in AGO2 crosslinking near annotated miRNA target sites relative to control cells. Taken together, these data suggest an attenuation in AGO2 binding on pro-proliferative mRNAs in presence of I3.

### IGF2BP3 alters the 3’-UTR occupancy of several tumor suppressive and tumorigenic microRNAs

Our data suggests antagonization of AGO2 by I3 on 3’-UTRs bound by both RBPs. To determine if I3 expression impacts the occupancy of all miRNAs or only specific miRNAs, we queried miR-eCLIP data for I3-dependent changes in chimeric read representation. We normalized chimeric read counts for all miRNAs bound to 3’-UTR sequences to total chimeric read counts in each condition obtained from miR-eCLIP. We found that miR-181a, miR-20a, miR-17 and miR-19b have significantly different chimeric read counts between both cell lines despite overall levels of chimeric reads being indistinguishable between cell lines (Supp. Fig.3A-B). Intriguingly, miR-181a showed a significantly higher 3’-UTR chimeric read count in the I3KO, while miR20a, miR17 and miR19b showed the opposite trend (Supp.Fig.3A). Small RNA sequencing confirmed that the steady-state levels for the majority of miRNAs, including miR181a, -20a, -17 and -19b, are not affected by I3 knock-out (Supp. Fig.3C). To further assess the observed chimeric read difference, we quantified the differential chimeric read occupancy (DCRO) for miR-181a, -20a, -17 and 19b on each of their respective target 3’-UTRs (Fig.3A) The miR-181 family and miR-142 have well established roles in modulation of hematopoiesis ^26,27^ and are among the most abundant miRNAs in our sRNA-seq data set and in our chimeric reads. For this reason, we included miR-181b and miR-142 in this analysis, along with let-7a and let-7f due to the known tumor-suppressive role of the let-7 family. We identified a total of 200 mRNA targets with DCRO at their 3’-UTRs for the selected miRNAs (Fig.3A). Significant increases in DCRO in the I3KO are marked in purple, while significant decreases in DCRO in the I3KO are marked in blue. We found that miR-181a exhibits the most instances of DCRO, all of which are significantly increased in the I3KO (Fig.4, Table 2). Figure 3B shows an example gene, PUM1, with this miR-181a binding pattern at its 3’-UTR. Interestingly for let-7a, let-7f, miR-20 and -19b we observe examples of both strongly increased and decreased DCRO, while miR-142 and -17b exclusively show decreased DCRO in the I3KO (Fig.3A, Table 2).

**Figure 3.**
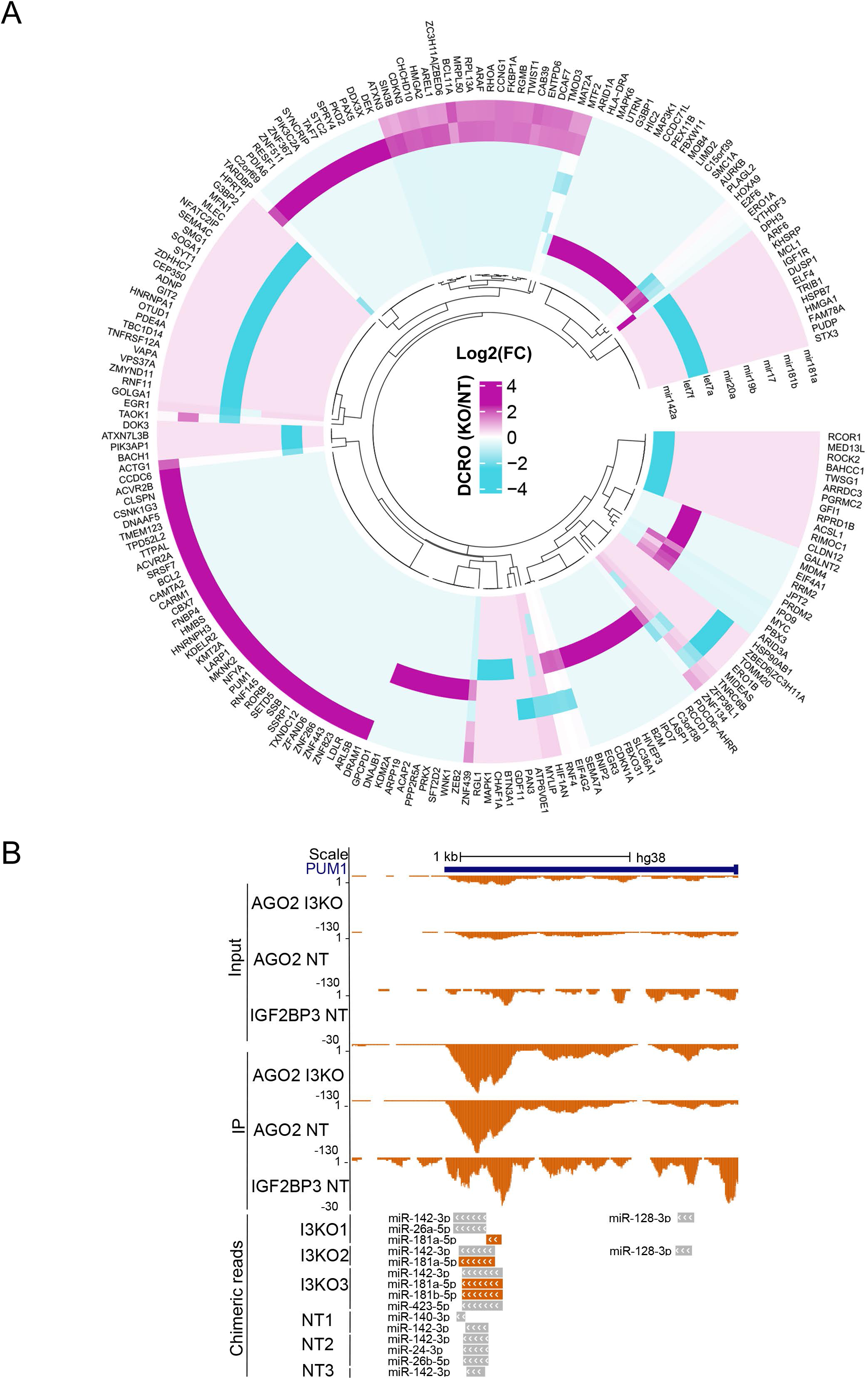
IGF2BP3 remodels the targeting landscape of cancer-related miRNAs. (A) Circle plot depicting Differential Chimeric Read Occupancy (DCRO) of miR-181a/b, miR-17, miR-19b, miR-20, let-7a, let-7f and miR-142 between control (NT) and I3 knock-out (I3KO) cells, with respect to I3KO with a log2FC of +1 and -1 and p value of 0.05. (B) UCSC genome browser screenshot showing AGO2 chimeric eCLIP peaks on the 3’-UTR of the I3-dependent miR-181 target *PUM1*. Non-chimeric AGO2 coverage tracks in control and I3 knock-out (I3KO), as well as I3 coverage tracks in control are shown as a reference. Size-matched inputs for each immunoprecipitation coverage track are also shown.

**Figure 4.**
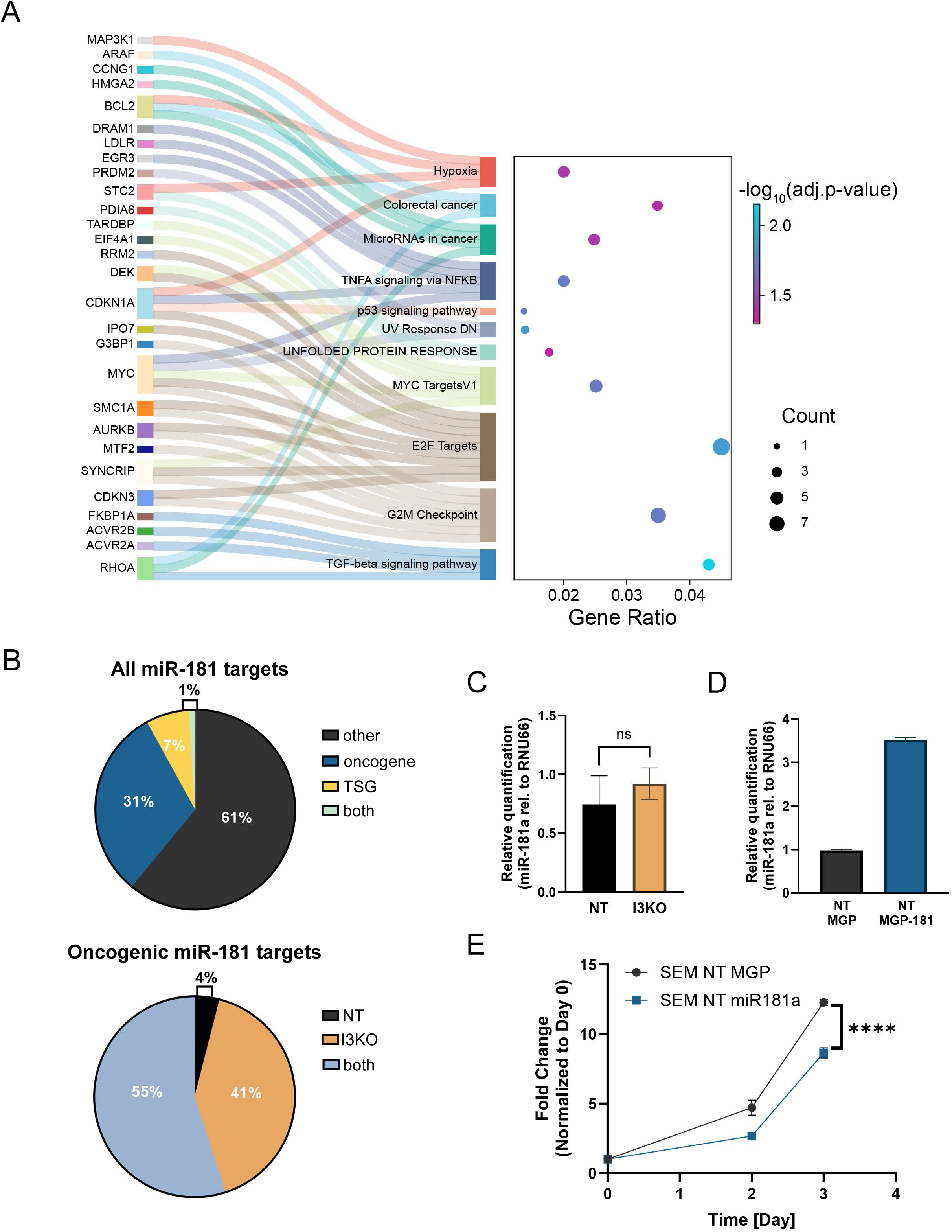
(continued) IGF2BP3 remodels the targeting landscape of cancer-related miRNAs. (A) Hallmark enrichment analysis of mRNAs with increased Differential Chimeric Read Occupancy (DCRO) from Figure 3. Significantly enriched pathways had an FDR of less than 0.05. Genes part of the enriched pathways are annotated and connected by a Sankey plot. (B) The proportion of miR-181 targets annotated as either oncogenes or tumor suppressors according to the TSGene database is shown on top. The proportion of oncogenic miR-181 targets from the left pie chart bound by miR-181 in either control (NT, black), I3 knock-out (I3KO, orange) or both (blue) is shown on the bottom. (C) Quantification of miR-181 levels in control (NT) and I3 knock-out (I3KO) SEM cells. RQ values of miR-181 relative to the small RNA control RNU66 are plotted. A paired t-test was used to calculate statistical significance. (D) Quantification of miR-181a in control cells with empty vector MGP (NT MGP, black), and control with miR-181 overexpression (NT MGP-181a, blue). RQ values normalized to the endogenous small RNA control RNU66 using the 2^−ΔΔCt^ method are plotted for each condition. (E) Quantification of cell growth using the CellTiter-Glo assay of SEM cells with empty vector MGP (NT MGP, black) and control with miR-181 overexpression (NT MGP-181a, blue). Fold-change of luminescence relative to day 0 is plotted for each condition. 2-way ANOVA; p-value <0.0001

**Table 1.**
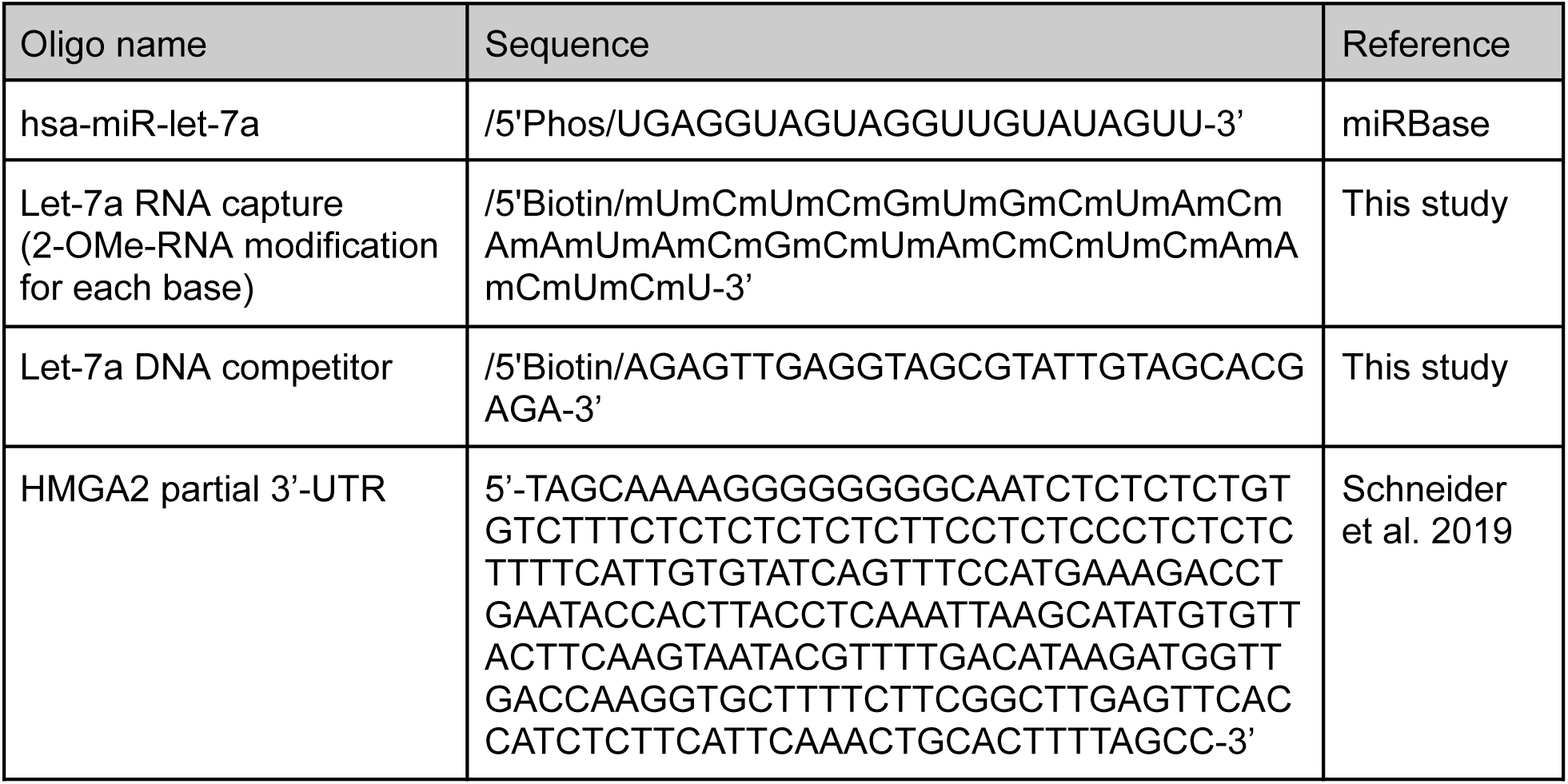
List of oligonucleotides used in this study:

**Table 2.**
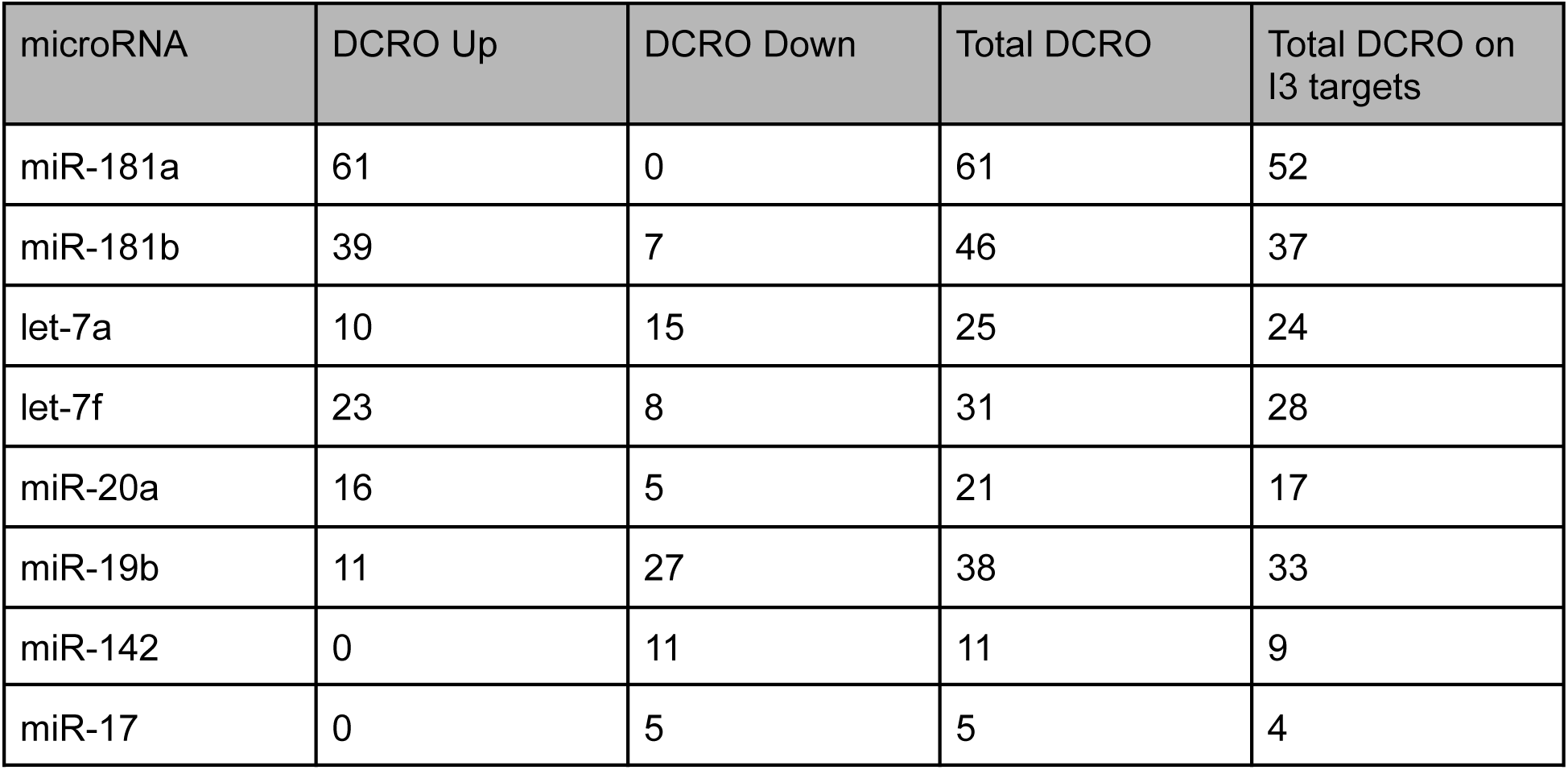
Number of Differential Chimeric Read Occupancy (DCRO) events for the selected microRNAs with respect to IGF2BP3 knock-out cells:

Supplementary figure 3E and F shows two example 3’-UTRs, SMC1A and SMG1, with increased let-7 DCRO and decreased miR-19b DCRO in the I3KO respectively. Hallmark enrichment analysis of all 200 mRNA targets in Figure 3A revealed that mRNAs with increased DCRO in the I3KO have an enrichment for cancer-related processes, including MYC-related genes (MYC-Targets V1), p53 signaling and regulation of cell cycle (E2F Targets, G2M Checkpoint) among others (Fig.4A). By contrast, there were no significantly enriched Hallmark processes for mRNAs with decreased DCRO in the I3KO. We identified a few significantly enriched KEGG pathways, albeit with low gene counts (Supp.Fig.3D). To determine if I3 has a direct effect on miRNA occupancy, we quantified the number of DCRO on mRNAs bound by I3 and observed a similar result than for the previous set of mRNAs (compare Supp.Fig.4A with Fig.3A). Enrichment analysis showed several enriched GO Biological processes belonging to transcriptional regulation, regulation of cell cycle and signaling pathways, among others (Supp.Fig.4B).

miR-181 is hypothesized to function as a tumor suppressor in several types of cancer^28^. In our data set, miR-181a exhibits the highest number of DCRO, all of which represent exclusively increased DCRO in the I3KO. To determine if I3 might be interfering with miR-181 targeting of oncogenic transcripts, we quantified changes in miR-181 occupancy in control or I3KO cell lines using our chimeric read set. We found that 31% of 3’-UTRs targeted by miR-181 in either I3KO or control cells correspond to oncogenes (Fig. 4B, left panel). Intriguingly, deletion of I3 results in a significant increase in miR-181-oncogene association (Fig 4B, right panel).

Previous studies from our group and others characterized a consistently decreased cell growth phenotype in I3KO cells compared to control across several different leukemia cell types ^10,11,13^. Our data shows that in absence of I3, miR-181 exhibits the most instances of significantly increased DCRO on cancer-related mRNAs (Fig.3A, Fig.4A, Table 2), increased association on oncogenic 3’-UTRs (Fig.4B), despite overall steady-state miR-181 levels being unaffected by the I3KO (Fig.4C, Supp.Fig.3B). These observations suggest a competitive relationship between miR-181 and I3. To test the hypothesis that I3 and miR-181 compete *in vivo*, we created SEM cell lines with miR-181a overexpression (Fig.4D). If I3 attenuates miR-181 activity, we expect overexpression of miR-181 to mimic the phenotype of I3 knock-out cells. We observed that SEM cells overexpressing miR-181 grew significantly slower than control SEM cells (Fig.4E). Taken together, our data suggest an I3-dependent alteration in miRNA occupancy on transcripts involved in cancer-related pathways, resulting in I3-dependent promotion of cell proliferation, and indicates an antagonistic relationship between I3 and miRNAs on common mRNA targets.

### IGF2BP3 directly competes with AGO2

Our previous results indicate an antagonization of tumor suppressive AGO2-bound miRNAs by I3. To study the molecular binding modality of AGO2 and I3 and to either exclude or support a competitive model, we created recombinant proteins for *in vitro* binding studies. Figure 5A summarizes the experimental setup for Electrophoretic Mobility Shift Assays (EMSA) as well as expected outcomes of the EMSA. Recombinant human I3 as well as human AGO2 loaded with a miRNA of choice were expressed in Sf9 insect cells and affinity purified using tags (Supp. Fig. 5A-B). AGO2 bound to a miRNA is known as “minimal RISC” and is in itself sufficient to carry out its function at the miRNA binding site^29,30^. The RNA target was transcribed *in vitro* and radioactively labeled at the 5’-end. As a proof of concept for our binding studies, we chose a 225bp sequence from the HMGA2 3’-UTR, which was already used successfully by Schneider and colleagues for EMSA with I3^14^. To expand on these assays, we performed EMSA of AGO2-miR complexes in the presence of I3 at varying concentrations (Fig. 5B). Since the HMGA2 substrate carries let-7 recognition sites, we generated recombinant AGO2-let-7. AGO2-let-7 was titrated in a two-fold dilution series ranging from 0.025nM to 2nM. As controls, shifts without any I3 protein (lane 1) and without AGO2 (lane 2,3) were performed. The middle band corresponding to AGO2 becomes fainter as the I3 concentration increases. This effect was reproducible across multiple experiments (n=3). The binding pattern visible on the EMSA suggests a competitive mode of binding for I3 and AGO2. Since the I3 recombinant protein was purified with its GST tag attached, we performed EMSA of I3 and the GST tag by itself on HMGA2 RNA, to exclude the possibility of the GST tag contributing to binding (Supp. Fig.5C). We confirmed that the GST tag by itself does not bind to our RNA substrate. Taken together, our data supports a competitive mode of interaction between I3 and RISC, in which I3 directly competes with AGO2 for miRNA binding sites (Fig.5C).

**Figure 5.**
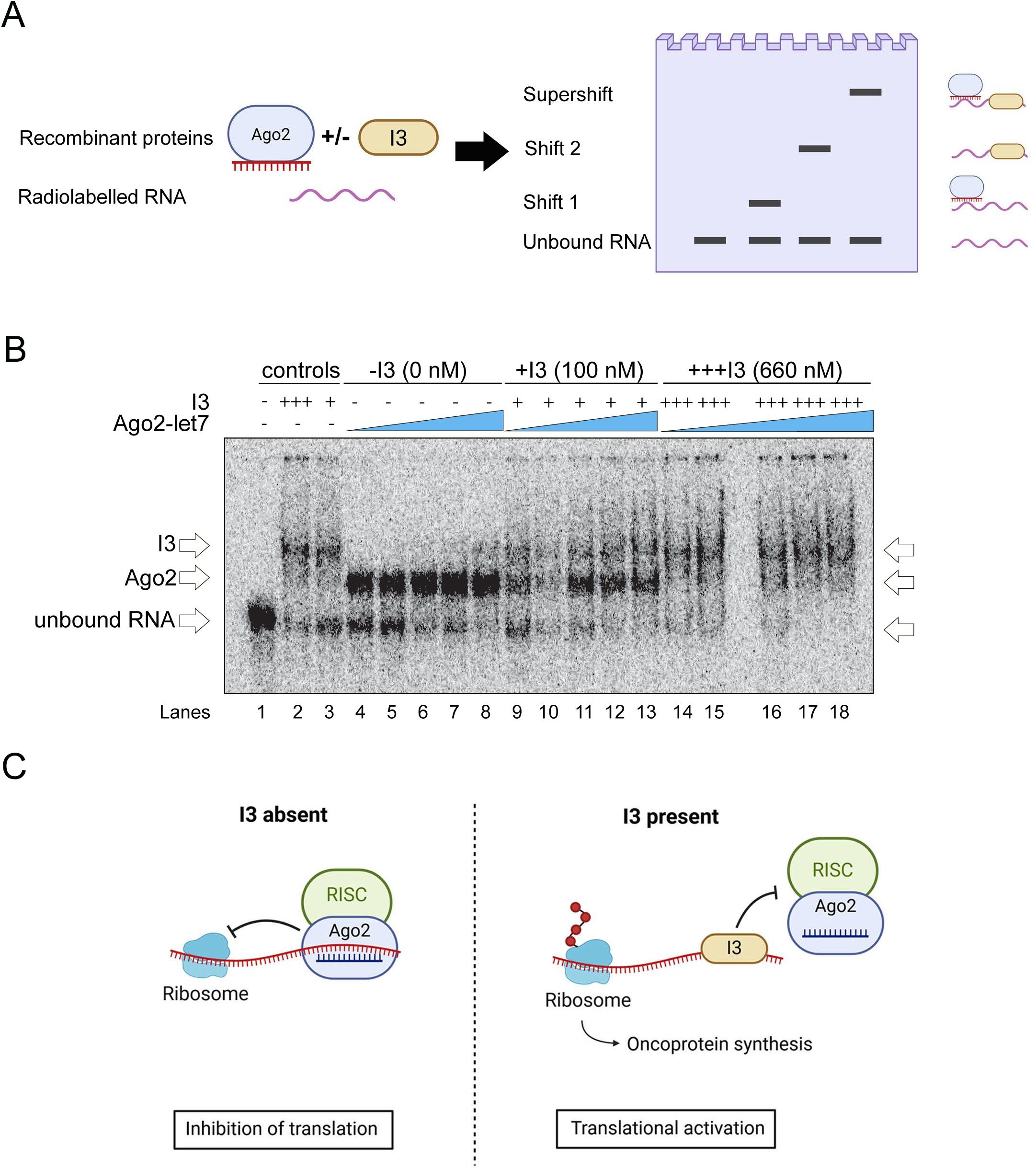
IGF2BP3 directly competes with AGO2 *in vitro*. (A) Schematic overview of the Electrophoretic Mobility Shift Assay (EMSA) experiment. I3 and AGO2 were expressed using the Baculovirus system and affinity purified using GST and Nickel sepharose resin respectively. For AGO2, the affinity purification was combined with loading of a synthetic microRNA. Target RNA was transcribed *in vitro* using T7 promoter on the pBluescript I KS (+) plasmid. The RNA was purified, radioactively labelled and subsequently used for EMSA with recombinant I3 and AGO2-miRNA complexes. (B) Electrophoretic Mobility Shift Assay (EMSA) using recombinant I3 and AGO2 loaded with synthetic let-7a microRNA. A partial 3’-UTR sequence from the oncogene HMGA was used as the RNA substrate. Lane 1 = no protein control, Lane 2 = saturating (660nM) I3 only, Lane 3 = sub-saturating (100nM) I3 only, Lane 4-8 = AGO2-let-7a titration without I3, Lane 9-13 =AGO2-let-7a titration with sub-saturating I3, Lane 14-18 = AGO2-let-7a titration with saturating I3. (C) Illustration of model for AGO2 and I3 binding in leukemic cells. In the absence of I3, RISC binds to its microRNA target site, inhibiting oncoprotein synthesis, thus downregulating tumorigenic genes. In the presence of I3, such as our leukemic SEM cell model, I3 binds to and occludes microRNA binding sites. This inhibits RISC function and favors translation of tumorigenic transcripts, thus amplifying cancer hallmarks.

## Discussion

Previous work from our group and others proposed that I3 modulates interactions between the RNA-induced silencing complex (RISC) and the 3′ untranslated regions (3′UTRs) of target mRNAs, thus shaping miRNA-mediated repression ^13,14,17^. Our findings support this model and refine it by demonstrating direct competition between I3 and RISC at target sites. Specifically, we show that I3 inhibits the binding of select cancer-associated miRNAs, including miR-181 and let-7, by directly competing with RISC for access to 3′UTR elements, promoting cellular pathways aligned with cancer hallmarks.

I3 is a multifunctional RNA-binding protein (RBP) involved in diverse post-transcriptional mechanisms ^15^. Earlier work suggested that I3 protects mRNAs from miRNA-mediated degradation by sequestering transcripts within membrane-less “safe house” compartments^18^. While our results are consistent with an overall antagonism of miRNA function, they diverge from a strictly sequestration-based model. Instead, we provide evidence that I3 actively interferes with miRNA–mRNA association by directly competing with RISC. At the global level, mapping I3 and I3-dependent AGO2 binding sites in MLL-AF4 B-lymphoblastic leukemia cells, with and without I3 expression, revealed that loss of I3 increases AGO2 occupancy on mRNAs involved in proliferation, including oncogenes such as *HOXA7*, *BCL2L11*, *MYB*, and *SOX4*.

These data indicate that I3 reduces RISC binding on oncogenic transcripts through a competitive 3′UTR engagement. Mechanistically, Electrophoretic Mobility Shift Assays (EMSAs) demonstrate that I3 directly competes with AGO2 at the *HMGA2* 3′UTR, displacing RISC *in vitro*. While *in vitro* systems lack the complexity of a cellular environment, together with the global occupancy shifts they strongly support a competitive relationship between I3 and RISC that constrains miRNA access to oncogenic targets.

Beyond global antagonism, our analyses uncover a bi-modal regulatory mode of action where I3 differentially mediates RISC binding in a miRNA- and context-specific manner. Quantification of differential miRNA occupancy across 3′-UTRs shows that I3 antagonizes RISC binding for miR-181 and let-7, facilitates RISC binding for miR-142 and miR-17b, and exhibits mixed effects for miR-20a, miR-19b, and let-7a/f. This pattern indicates that I3 does not uniformly block miRNA function but rather fine-tunes targeting to reinforce a malignant gene-expression network. Functionally, miR-181 displays context-dependent roles in hematologic malignancies^31–33^. Although it has been linked to both tumor-suppressive and oncogenic mechanisms, our data in human leukemic B cells indicate that I3 antagonizes miR-181 binding on oncogenic transcripts. By contrast, let-7 is a well-established tumor suppressor family, whereas miR-20a, miR-19b, and miR-17b are oncogenic miRNAs encoded within the same cluster ^34–38^. The mixed I3-dependent effects we observe across these miRNAs are consistent with a selective, transcript- and miRNA-specific mode of I3 action. Overexpression of miR-181a in I3 knock-out cells rescues the known growth defect, consistent with a model in which increasing miRNA levels can overcome I3-mediated antagonism. Taken together, these results recapitulate and extend our previous proposal that I3 plays a bi-modal role, inhibiting RISC binding on oncogenes while facilitating RISC association on tumor suppressors^17^. One plausible mechanism underlying I3’s dual roles is RNA structural remodeling. I3 binds broadly across 3′UTRs, occupying large RNA footprints with low sequence specificity^14^, and could thereby alter local RNA conformation to either expose or occlude miRNA target sites. Because RISC activity is sensitive to RNA secondary structure ^39^, we hypothesize that I3-dependent modulation of RNA folding fine-tunes miRNA accessibility in a site- and miRNA-specific manner. This hypothesis suggests multiple directions for future work. First, miR-eCLIP can be miRNA-enriched (as in murine T-cells by Verheyden and colleagues ^40^) or read-sub-sampled (as in Hofmann et al. ^25^) to dissect the broader interactome of individual miRNAs in the presence or absence of I3. Second, intrinsic miRNA–site affinity may contribute to I3’s remodeling effects ^41^; stratifying targets by predicted/empirical affinity and local structure could clarify when I3 acts as an antagonist versus a facilitator.

At the systems level, the global shift to a stem-like gene-expression program observed in this study and previously likely arises from a combination of miRNA-binding antagonism/remodeling along with direct and indirect effects of I3 on its RNA partners. One direct mechanism is mRNA stabilization, reported in acute myeloid leukemia ^12^. Consistent with stabilization, we observe a significant enrichment of I3 mRNA targets among differentially expressed transcripts. In parallel, the overrepresentation of I3 targets among differentially expressed proteins in our dataset suggests an additional role in translational regulation. Such translational control by I3 has been documented both during development (physiological I3 expression) and in cancer contexts (aberrant I3 expression) ^18,22,42^ ^20^. These convergent mechanisms position I3 as a node of post-transcriptional control that coordinates transcript stability, structure, and translation to promote leukemogenic programs.

In summary, our study identifies I3 as a versatile regulator of miRNA function that directly competes with RISC for 3′-UTR access, bi-modally modulates miRNA activity in a miRNA- and context-specific manner, and remodels post-transcriptional gene regulation through effects on RNA structure, mRNA stabilization, and translation. By fine-tuning access of oncogenic and tumor-suppressive miRNAs to their targets, I3 emerges as a key determinant of malignant gene-expression programs in leukemia. These insights broaden our understanding of post-transcriptional regulation in hematologic malignancies and highlight I3 as a promising therapeutic target and biomarker.

## Materials and Methods

### Cell culture

The human B-ALL cell line SEM (DMZ-ACC 546) was cultured as previously described^43^. All cell lines were maintained in standard conditions in an incubator at 37 °C and 5% CO_2_ and cultured in IMDM with 10% FBS.

### CRISPR/Cas9-mediated deletion of IGF2BP3 in SEM cells

An IGF2BP3 deletion (I3KO) in SEM cells was introduced using the lentiviral delivery of CRISPR/Cas9 components in a two-vector system and sgRNA sequence as previously described^10,43^.

### Total RNA extraction

SEM cells were pelleted by centrifugation at 200 x g for 5 minutes and resuspended in TRIzol (Invitrogen). Samples were stored at -80C until use. Total RNA was extracted using the TRIzol protocol (Invitrogen) for isolation of RNA from cells in suspension. After the total RNA was solubilized in ddH_2_0, one overnight ethanol precipitation step was included for further purification of the total RNA.

### eCLIP sequencing of I3 and AGO2

IGF2BP3 eCLIP and AGO2 miR-eCLIP studies were performed by Eclipse Bioinnovations Inc (https://eclipsebio.com/) according to the published eCLIP and miR-eCLIP protocols ^24,44^. SEM cells were grown and UV cross-linked at 400 mJoules/cm^2^ with 254 nm radiation, flash frozen, and stored until use at −80°C. Crosslinked cell pellets were further processed by Eclipse Bioinnovations for eCLIP using a validated rabbit anti-IGF2BP3 antibody (MBL, RN009P) and Eclipse AGO2 antibody (Santa Cruz Biotech sc-53521). For IGF2BP3, three replicates using 20 million SEM NT cells per replicate and one HEPG2 control were processed, yielding a total of eight libraries (4 IP libraries and 4 size-matched input libraries). For the AGO2 miR-eCLIP data set, 6 IP replicate libraries were generated using 17 million cells per replicate: 3 from SEM NT and 3 from SEM I3KO. 6 size matched inputs (SMI) were generated. Sequencing was performed as SE72 on the NextSeq platform.

### eCLIP read processing

Reads from IGF2BP3 and AGO2 eCLIP were pre-processed for quality and adapter trimming (cutadapt; RRID:SCR_011841) followed by alignment to the reference human genome (GRCh38, GENCODE v41 annotation) using STAR v2.7.7a. Repetitive elements were removed by a reverse intersection of all peak files with the repeat masker bed file (downloaded from the University of California Santa Cruz [UCSC] table viewer; RRID:SCR_005780) while PCR duplicate reads were removed, and the aligned files were further processed and analyzed for peaks enriched over the background using Skipper v1.0.087 (RRID:SCR_026260). IGF2BP3 and AGO eCLIP fine-mapped peak sets were filtered for peaks with log2(fold-change) ≥ 1.0 and ≥ 3.0, respectively, in terms of mean read counts in IP vs. size-matched input. Peaks were annotated using transcript information from GENCODE v41 with the following priority hierarchy to define the annotation of overlapping features: protein coding transcript (CDS, UTRs, intron), followed by non-coding transcripts (exon, intron).

Differential binding in AGO2 I3KO and NT samples were analyzed using DESeq2 -1.32.0. (RRID:SCR_015687). Count files for each of the replicates were subjected to differential binding to identify statistically significant differences between I3KO and NT. The p-values from the Wald tests are displayed as the y-axis of volcano plots, with the log2 fold-change of normalized peaks intensities (I3KO over NT) as the x-axis.

Bedtools intersect module was used to find the peaks exclusive to NT and I3KO along with the overlapping peaks. Furthermore, the peak counts were normalized to CPM (counts per million) and used for visualization of peak intensities across CDS and UTRs for AGO NT and KO replicates. Python 3 was used for visualization of metagene and CDF plots while the ggplots package from R was used to plot gene enrichment.

### Chimeric AGO2 miR-eCLIP read processing

After sequencing, samples were processed with Eclipsebio’s proprietary analysis pipeline (v1). UMIs were pruned from read sequences using umi_tools (v1.1.1) followed by trimming of 3’ adapters from reads using cutadapt. Reads mapped to a custom database of repetitive elements and rRNA sequences were removed and remaining non-repeat mapped reads were mapped to the hg38 genome using STAR (v2.7.7a). PCR duplicates were removed using umi_tools (v1.1.1). AGO2 eCLIP peaks were identified within eCLIP samples using the peak caller CLIPper (v2.0.1). For each peak, IP versus input fold enrichments and p-values were calculated and mapped to repetitive elements or the genome using bowtie (v1.2.3). The miRNA portion of each read was then trimmed, and the remainder of the read was mapped to the genome using STAR (v2.7.7a). PCR duplicates were resolved using umi_tools (v1.1.1), and miRNA target clusters were identified using CLIPper (v2.0.1). Each cluster was annotated with the names of miRNAs associated with that target. Peaks were annotated using transcript information from GENCODE v41 to define the final annotation of overlapping features: protein coding transcript (CDS, UTRs, intron), followed by non-coding transcripts (exon, intron). Further, the top differentially expressed miRNAs were selected from small RNA sequencing data and their chimeric read proportions were calculated in NT and I3KO and plotted using GraphPad Prism.

To limit our analysis to biologically relevant AGO2 peaks, chimeric AGO2 peaks were overlapped with the total AGO2 peaks from the eCLIP dataset. AGO2 peaks present in both data sets were further used for downstream analysis.

### Quantification of Differential Chimeric Read Occupancy (DCRO)

miRNAs exhibiting differences in chimeric read proportions between NT and I3KO were further analyzed to assess the chimeric read occupancy at the mRNA target level for each miRNA. Chimeric peak counts corresponding to the annotated 3’ UTR targets for each miRNA were considered individually for NT and I3KO replicates to calculate the differential chimeric read occupancy. DEseq2 was used to identify the significant differential chimeric read occupancy across mRNA targets of each miRNA. A matrix of miRNA targets and their corresponding chimeric read occupancy in terms of log2 fold changes was generated and visualized as a circular heatmap using *ComplexHeatmap* and *circlize* packages in R.

### HOMER motif analysis

The peak bed files from AGO2 miR-eCLIP and IGF2BP3 eCLIP were utilized to identify known and *de novo* motif sequence enrichment for NT and I3KO using findMotifsGenome.pl from Homer2 with default and modified parameters for RNA motifs with a threshold length of 10.

### Small RNA sequencing

Total RNA from SEM NT and I3KO cells were used for sRNA library preparation. 1 μg of total RNA for each sample was used for sRNA library preparation using the NEXTFLEX small RNA-Seq Kit v3 following the manufacturer’s protocol (Perkin Elmer Applied Genomics). Pooled sRNA sequencing libraries were sequenced on an Illumina HiSeq 4000 at the UC Davis Sequencing Core Facility, generating 100 bp single-end reads.

### mRNA sequencing

Total RNA from SEM NT and I3KO cells was used for mRNA library preparation. 2 μg of total RNA for each sample was used for mRNA library preparation using the NEXTFLEX Rapid Directional RNA-Seq Kit 2.0 following the manufacturer’s protocol (Perkin Elmer Applied Genomics). Before library preparation, total RNA samples were subjected to Poly(A) selection and purification using the NEXTFLEX Poly(A) Beads Kit 2.0 following the manufacturer’s protocol (Perkin Elmer Applied Genomics). Pooled mRNA sequencing libraries were sequenced on an Illumina NovaSeq S4 at the UC Davis Sequencing Core Facility, generating 150 bp paired-end reads.

### Proteomics analysis

50 μg of protein lysates were reduced, alkylated, and digested with trypsin using protein aggregation capture (PAC) as described in Batth et al^45^. Digested peptide samples were analyzed by LC-MS/MS on a Vanquish Neo UHPLC system coupled to a Thermofisher Orbitrap Astral mass spectrometer. Data was acquired using a data-independent analysis (DIA) acquisition strategy as described^46^. Data analysis was performed using the DIANN algorithm for peptide and protein identification and quantitation followed by Fragpipe analyst for comparison testing^47,48^.

### Generation of miR-181a overexpression cell lines

For the generation of SEM miR-181a overexpression cell lines, the full sequence of miR-18a (5’-AACAUUCAACGCUGUCGGUGAGU-3’) was cloned into the modified retroviral pMSCV vector MGP (Addgene plasmid # 26526; http://n2t.net/addgene:26526 ; RRID:Addgene_26526). The retroviruses were generated using pGAG/POL helper plasmid and pseudotyped with pVSVG. IGF2BP3 control cells were then stably transduced with retroviral supernatant overexpressing miR-181a and selected with puromycin.

### Quantification of miR-181a by qPCR

miR-181a and RNU66 levels were quantified in SEM cell lines using the TaqMan™ MicroRNA Assay (Applied Biosystems) according to the manufacturer’s instructions (assay ID 000480 and 001002 respectively).

### Quantification of cell growth using CellTiter-Glo

Empty MGP vector control or miR-181a overexpressing NT control cells were seeded at 1,500 cells/well in 96-well plates and cultured for 72 hours at 5% CO2 and 37°C. Cell growth was quantified using the CellTiter-Glo (Promega) assay according to the manufacturer’s instructions. Luminescence was measured using a Varioskan LUX multimode microplate reader (ThermoFisher). Five technical replicates were prepared for each sample.

### Human IGF2BP3 expression and purification

Recombinant IGF2BP3 tagged with GST was created using the Baculovirus protein expression system. Plasmid, Bacmid and P2 Baculovirus generation were carried out by Genscript. Subsequently, P2 virus was used to infect Sf9 insect cells. Sf9 cells were infected for 3 days at 27 °C and harvested by centrifugation. Cell pellets were resuspended in lysis buffer (20mM Tris-HCl pH 7.8, 100mM KCl, 0.2mM EDTA, 1mM DTT, 20% glycerol) and lysed in two passages through an Emulsiflex homogenizer (Avestin). Lysate was sonicated, clarified by centrifugation and tagged proteins were purified from the supernatant by incubating with Glutathione Sepharose 4B resin (Cytiva) for 20 minutes at 4°C. Resin was washed with Lysis Buffer and flow-through was continuously assessed for presence of contaminating protein by Bradford assay. After confirming that no more non-specific protein was present in the washes, IGF2BP3 was eluted off the resin using elution buffer (20mM Tris-HCl pH 7.8, 100mM KCl, 0.2mM EDTA, 1mM DTT, 20% glycerol, 10mM L-Glutathione). Elution was continuously assessed for presence of IGF2BP3 by Bradford until fully eluted from the resin. Eluted proteins were concentrated using a 50 kDa membrane Millipore concentrator. Subsequently, they were purified using size exclusion chromatography on a Superdex 200 Increase 10/300 GL column (Cytiva) on an Äkta Pure FPLC (Cytiva) using 20mM Tris-HCl pH 8, 100mM KCl, 0.2mM EDTA, 1mM DTT and 20% Glycerol. Protein concentration was determined by Bradford. Single-use aliquots were flash-frozen and stored at −80 °C until use.

### Preparation of human AGO2 loaded with synthetic miRNA

AGO2 loading with synthetic miRNA was combined with affinity purification and performed according to a previously published protocol with modifications^49^. Sf9 cells were infected for 72 hours at 27°C and harvested by centrifugation. Pelleted cells were resuspended in Lysis buffer (50mM NaH_2_PO_4_ pH 8, 0.3M NaCl, and 0.5mM TCEP) and lysed by two passages through an Emulsiflex homogenizer (Avestin). The lysate was clarified by centrifugation and Nickel affinity purification of AGO2 was performed using Ni Sepharose excel resin (Cytiva) for 1.5 hours at 4°C. Non-specific proteins were removed by washing the resin twice with 10 column volumes of wash buffer (50mM Tris pH 8.0, 0.3M NaCl, 15mM Imidazole, 0.5mM TCEP). Subsequently, the resin was washed with 10 column volumes of Calcium wash buffer (50mM Tris pH 8.0, 0.3M NaCl, 15mM Imidazole, 0.5mM TCEP, 5mM CaCl_2_). After the washes, a micrococcal nuclease treatment was performed to remove residual RNA. To that end, the resin was resuspended in Calcium Wash buffer and a micrococcal nuclease (Takara) was added. The resin was incubated for 40 minutes at room temperature with gentle inversion 2-3 times throughout the incubation period. After the digest, the resin was washed twice with 10 column volumes of Wash Buffer. AGO2 was eluted from the resin by resuspending it in 2 column volumes of Elution Buffer and incubating it on ice for 5 minutes. Eluted AGO2 was collected through gravity flow on a polypropylene column. The resin was resuspended once more in 1 column volume of the Elution buffer and flow-through was collected. EGTA was added to the purified AGO2 eluate to a final concentration of 5mM. Synthetic miRNA (Genscript) was then added to the eluate and incubated for approximately 5 minutes at 4°C. The resulting AGO2-miRNA mixture was dialyzed overnight at 4°C into Wash Buffer using a Slide-A-Lyzer 10kDa dialysis cassette (Thermo Scientific). Subsequently, AGO2 molecules loaded with the synthetic miRNA were purified and isolated from those bound to co-purifying cellular small RNAs as described previously ^49,50^.

Briefly, AGO2 loaded with a specific miRNA was captured using an antisense oligonucleotide (GenScript) and eluted with a DNA competitor (GenScript). Finally, the protein was dialyzed sequentially in Q Dialysis buffer (30mM Tris pH 8.0, 0.15M NaCl, 0.02% CHAPS, 0.5mM TCEP) and passed through Q Sepharose Fast Flow resin (Cytiva) to remove excess nucleic acids. Protein concentration was determined using Bradford. Single-use aliquots were flash frozen and stored at −80 °C until use.

The bacterial expression construct for full-length 6xHis-tagged human AGO2 protein was obtained from the lab of Professor Ian McRae (Scripps Research, San Diego, USA). A list of the used oligonucleotides can be found in Table 2.

### *In vitro* transcription and radioactive labeling of RNA substrates for Electrophoretic Mobility Shift Assay

The DNA sequence for the EMSA target RNA substrate (Table 2) was cloned and expressed using the pBluescript I KS (+) vector. *In vitro* transcription was performed in the presence of radioactive [α-32] UTP using the MEGAscript T7 High Yield Transcription Kit (Invitrogen) according to the manufacturer’s instructions. After precipitation, the transcribed RNA was run on a 5% Polyacrylamide gel, extracted from the gel and purified. Radioactive labeling efficiency was determined using a Beckman Coulter LS 6500 Multi-Purpose Scintillation Counter.

### Electrophoretic Mobility Shift Assay (EMSA)

EMSA was performed as previously described in CSH Protocols according to the “Simple Interactions” section ^51^.

### Data deposition

mRNA sequencing of NT and I3KO cells, as well as IGF2BP3 eCLIP were previously published^20^ and can be accessed under the BioProject accession number PRJNA1191523. Small RNA sequencing and AGO2 eCLIP data generated for this study can be accessed under the BioProject accession number PRJNA1339044. Liquid Chromatography-Tandem Mass Spectrometry data generated for this study can be accessed under the accession number MSV000099159 in the MASSive repository.

## Acknowledgements

We thank current and former members of the Sanford and Rao lab for their helpful discussions. We thank Professor Ian MacRae and Doctor Luca Gebert from the Scripps Research Institute for their support in optimizing the AGO2 guide loading protocol and for providing the AGO2 recombinant expression plasmid as well as Baculovirus stocks. We also thank Professor Seth Rubin at UC Santa Cruz and members of his lab for their helpful guidance in generating recombinant proteins for this study.

Research reported here was supported by the National Institutes of Health (NIH) and Eunice Kennedy Shriver National Institute of Child Health and Human Development (NIHCD) under award number T32HD108079 and the Institute for the Biology of Stem Cells (IBSC) at UC Santa Cruz. The content is solely the responsibility of the authors and does not necessarily represent the official views of the NIH/NICHD or the IBSC.

## Supplementary Figure Legends

**Supplementary figure 1.**
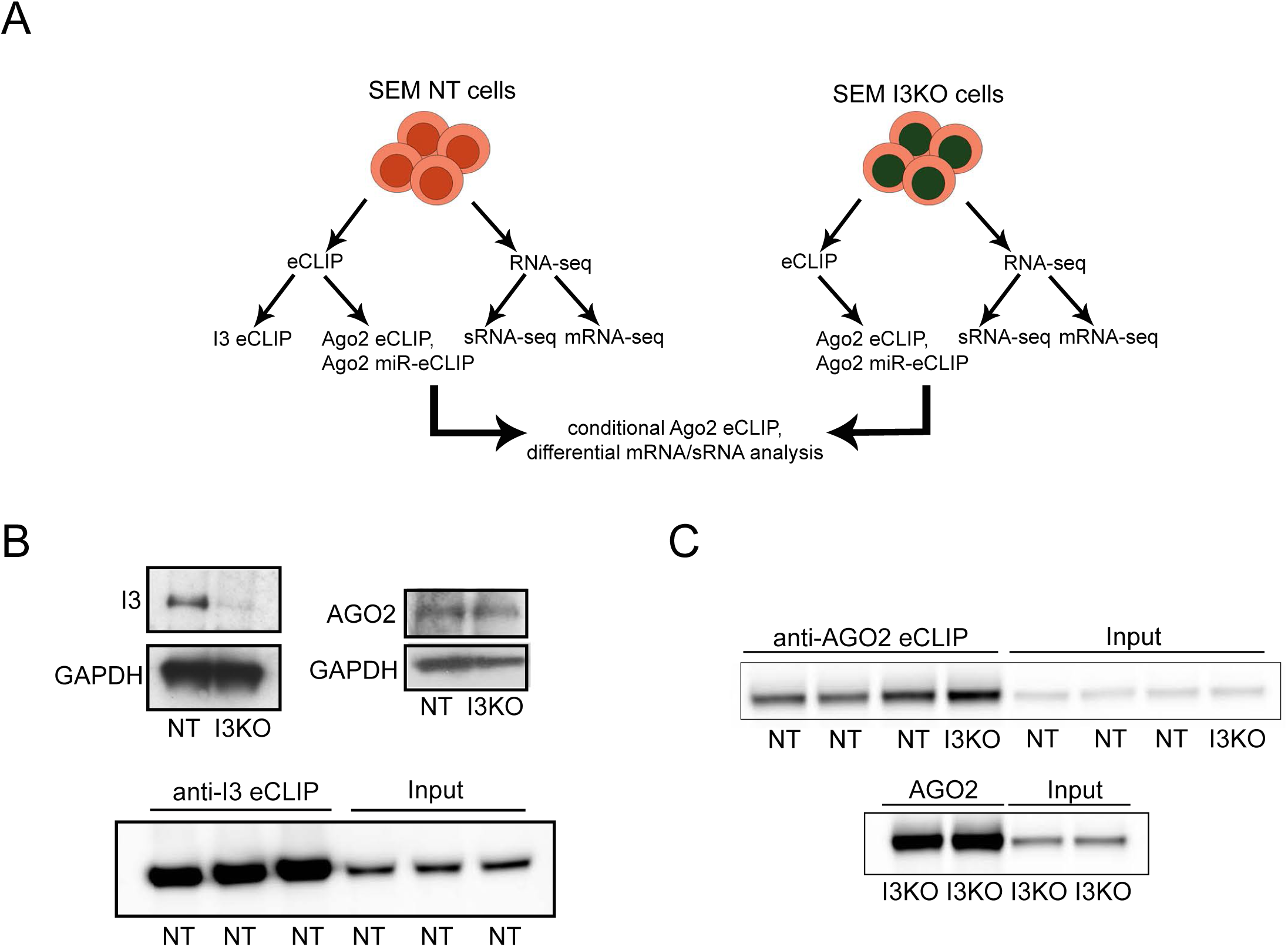
(A) Schematic overview of the cell lines and genomic sequencing data sets produced for this study. (B) Top: Western blots of control (NT) and I3-depleted (I3KO) SEM cells using antibodies against I3 and AGO2. GAPDH was used as a loading control. Bottom: Western blots of control (NT) SEM cells during I3 eCLIP using an antibody against I3. 10% of IP samples and 1% of input samples were loaded. (C) Western blots of control (NT) and I3-depleted (I3KO) SEM cells during AGO2 miR-eCLIP using an antibody against AGO2. 10% of IP samples and 1% of input samples were loaded and samples were run on two gels.

**Supplementary figure 2.**
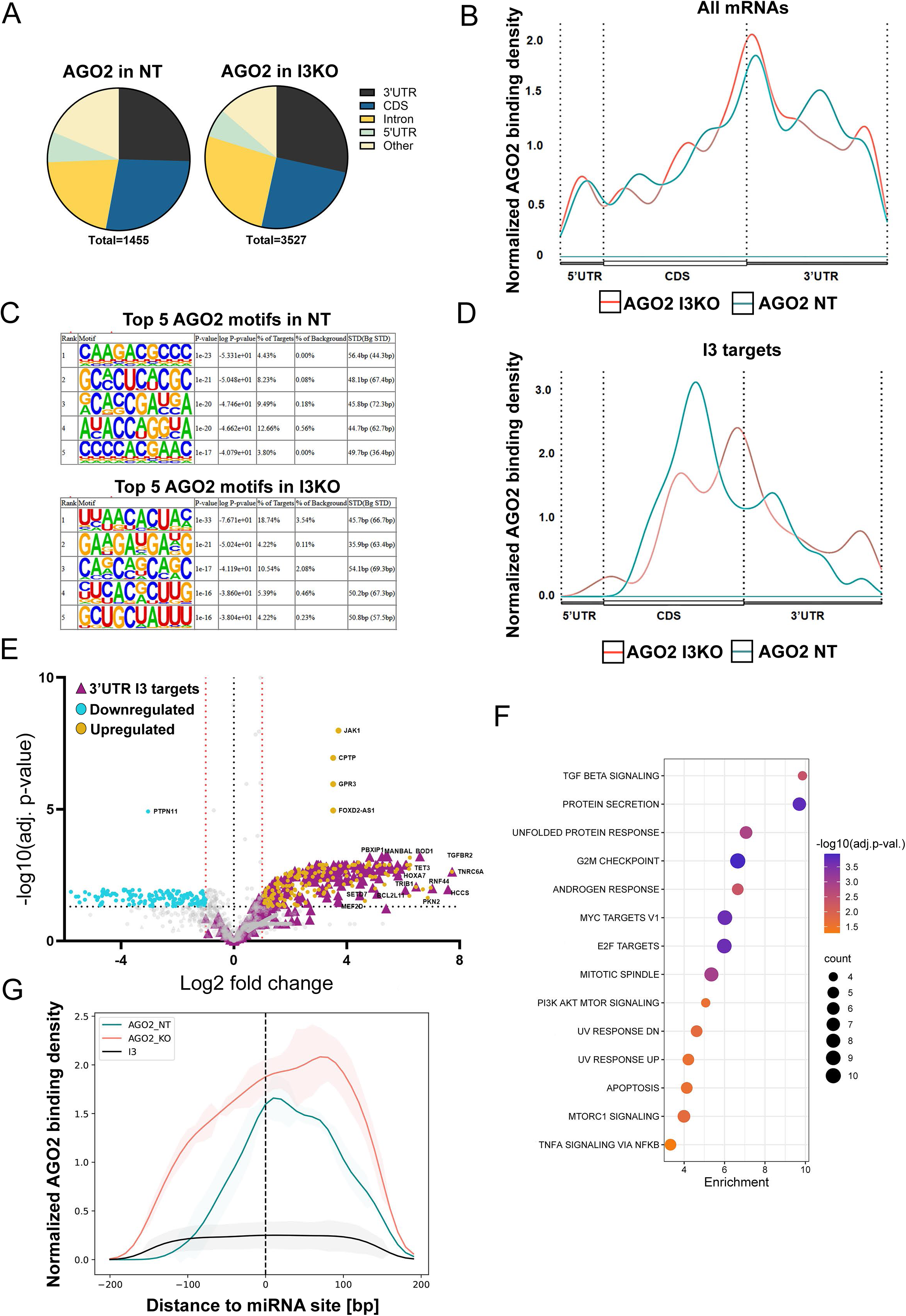
(A) Distribution of AGO2 eCLIP peaks across gene features in control (NT) and I3 knock-out (I3KO) SEM cells. Total number of peaks are written out for each condition underneath the respective pie chart. (B) Metagene plot depicting the relative density of AGO2 eCLIP peaks across gene features of all mRNAs in control (NT, blue) and I3-depleted (I3KO, red) SEM cells. Peak counts were normalized as counts per million relative to gene length. (C) Top five HOMER binding motifs of AGO2 eCLIP peaks in control (NT) and I3-depleted SEM cells (I3KO). (D) Same as B but with I3-bound mRNAs only. (E) Volcano plot showing differential AGO2 eCLIP peaks between control (NT) and I3-depleted (I3KO) cells, with respect to I3KO. Significantly different eCLIP peaks are depicted as colored dots. AGO2 eCLIP peaks in 3’-UTRs also bound by I3 in NT are marked as purple triangles. The threshold for differential peaks was defined as less than -1 Log2 fold-change or greater than 1 Log2 fold-change, with the p-value <0.05. (F) Hallmark enrichment analysis of mRNAs with increased 3’-UTR AGO2 eCLIP peaks in I3 knock-out cells (purple triangles from D). Significantly enriched pathways had an FDR of less than 0.05. (G) Metagene plot of AGO2 miR-eCLIP peaks in control (NT, blue) and I3 knock-out (I3KO, red) relative to annotated miRNA sites by miRBase. I3 eCLIP peaks in NT are depicted in grey. Peaks were normalized to CPM (counts per million) for all the replicates.

**Extended supplementary figure 2.**
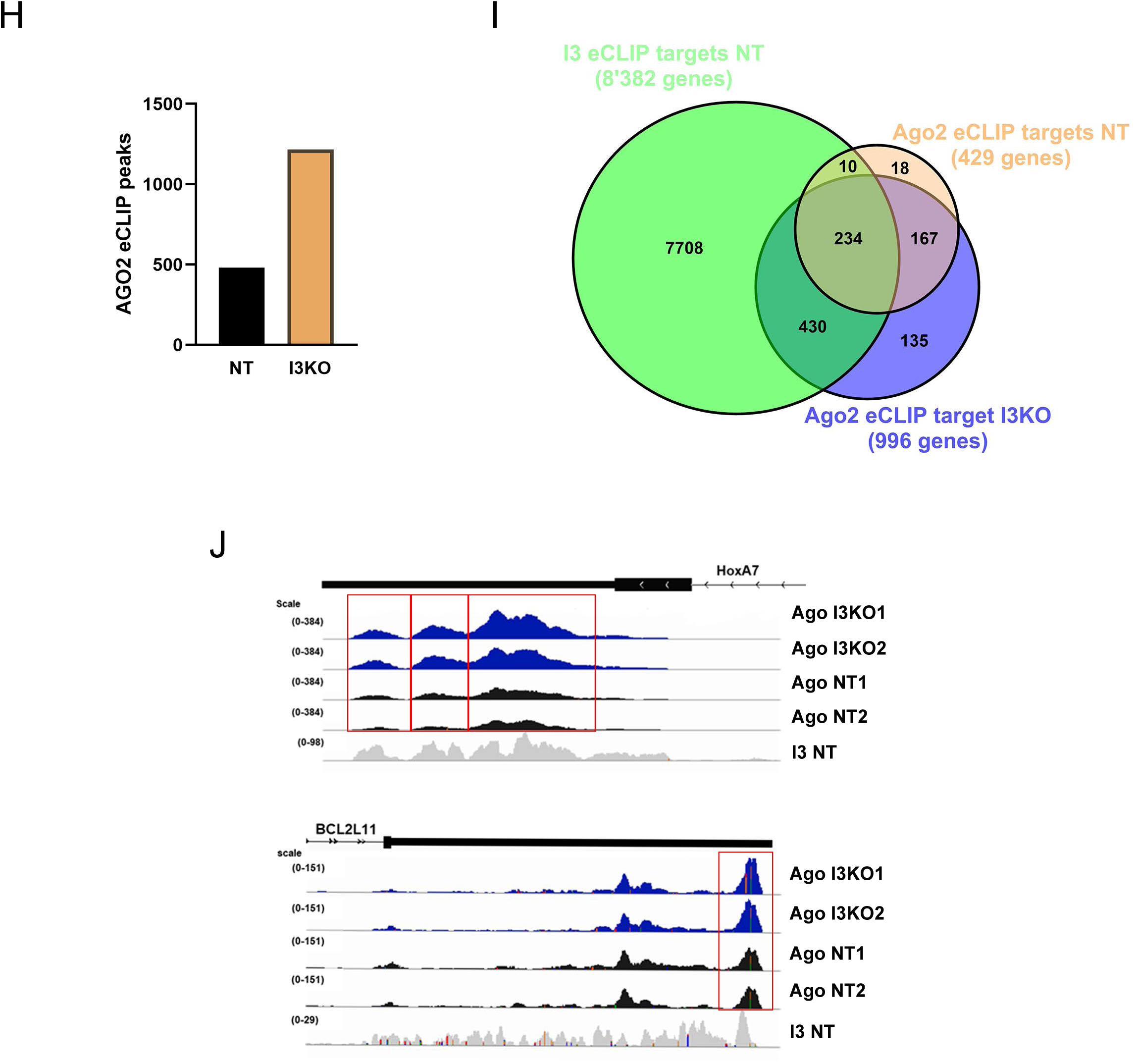
(H) Number of AGO2 eCLIP peaks in control (NT) and I3-depleted (I3KO) SEM cells. The cumulative number of peaks in each condition is plotted as a bar graph. (I) Venn diagram depicting the overlap of genes with eCLIP peaks from the I3 NT, AGO2 NT and AGO2 I3KO data sets. (J) IGV browser screenshots of *HOXA7* and *BCL2L11* 3’-UTRs. AGO2 NT and I3KO eCLIP tracks are shown in black and blue respectively. I3 NT eCLIP tracks are shown in grey. Differential AGO2 peaks are highlighted with red boxes. The eCLIP peaks were normalized to input using bamcompare and the resulting normalized files were plotted on IGV.

**Supplementary figure 3.**
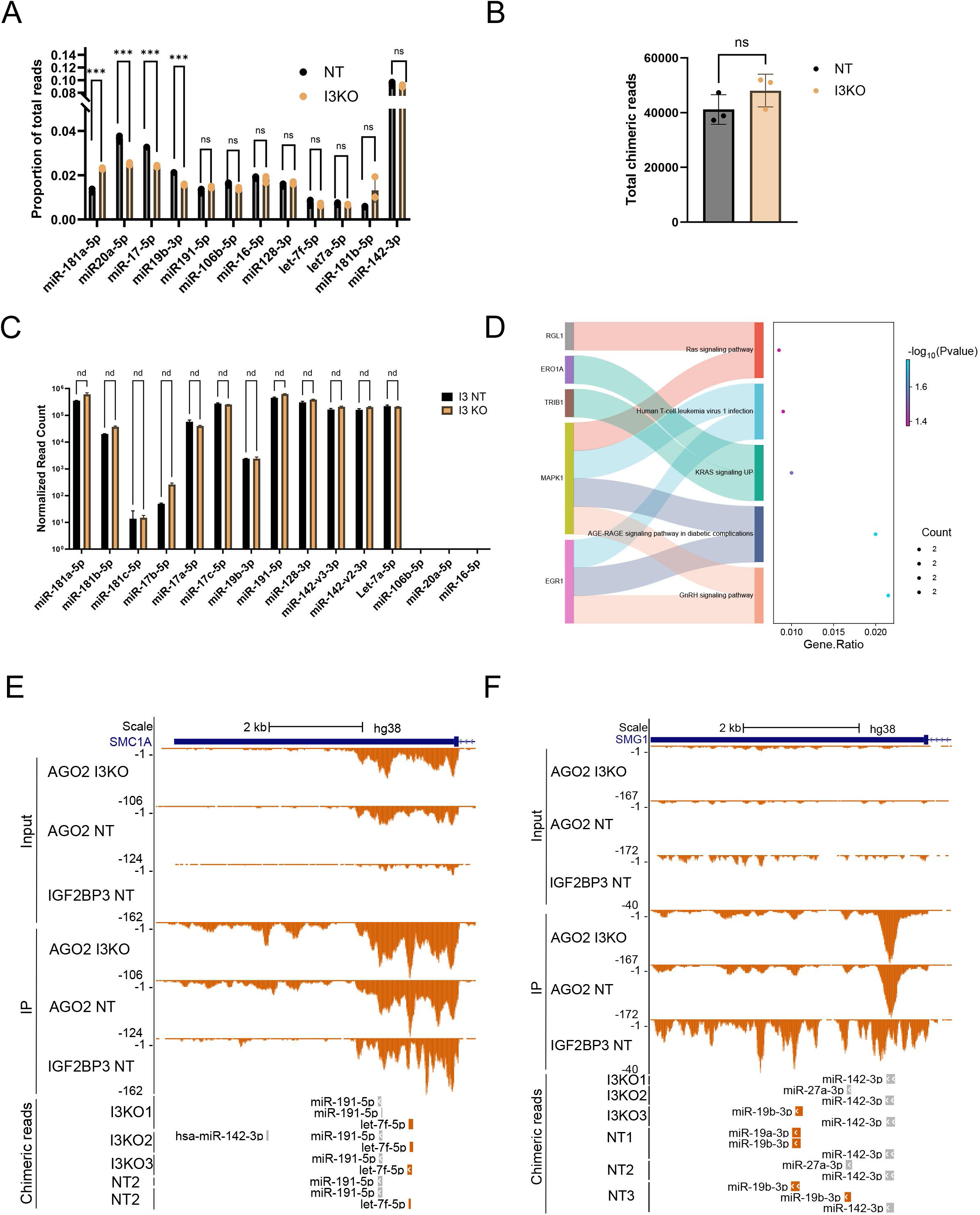
(A) Chimeric read count proportions of most abundant miRNAs mapping to 3’-UTRs from AGO2 miR-eCLIP reads in control (NT, black) and I3 knock-out (I3KO, orange) cells. Chimeric read counts for each miRNA were normalized to total chimeric read count in each condition, yielding read proportions. Two-way ANOVA was used to calculate statistical significance. (B) Quantification of total AGO2 miR-eCLIP chimeric reads across replicates in control (NT) and I3 knock-out (I3KO) SEM cells. An unpaired t-test was used to calculate statistical significance. (C) Normalized read counts of miRNAs identified by small RNA-seq in the control (NT, black) and I3 knock-out (I3KO, orange) cell lines. Two-way ANOVA was used to calculate the statistical significance. (D) KEGG pathway enrichment analysis of mRNAs with decreased Differential Chimeric Read Occupancy (DCRO) from figure 3A (E) UCSC genome browser screenshot showing AGO2 chimeric eCLIP peaks on the 3’-UTR of the I3-dependent let-7f target *SMCA1*. Non-chimeric AGO2 coverage tracks in control and I3 knock-out (I3KO), as well as I3 coverage tracks in control are shown as a reference. Size-matched inputs for each immunoprecipitation coverage track are also shown. (F) Same as E but showing the *SMG1* 3’-UTR with I3-dependent miR-19b binding.

**Supplementary figure 4.**
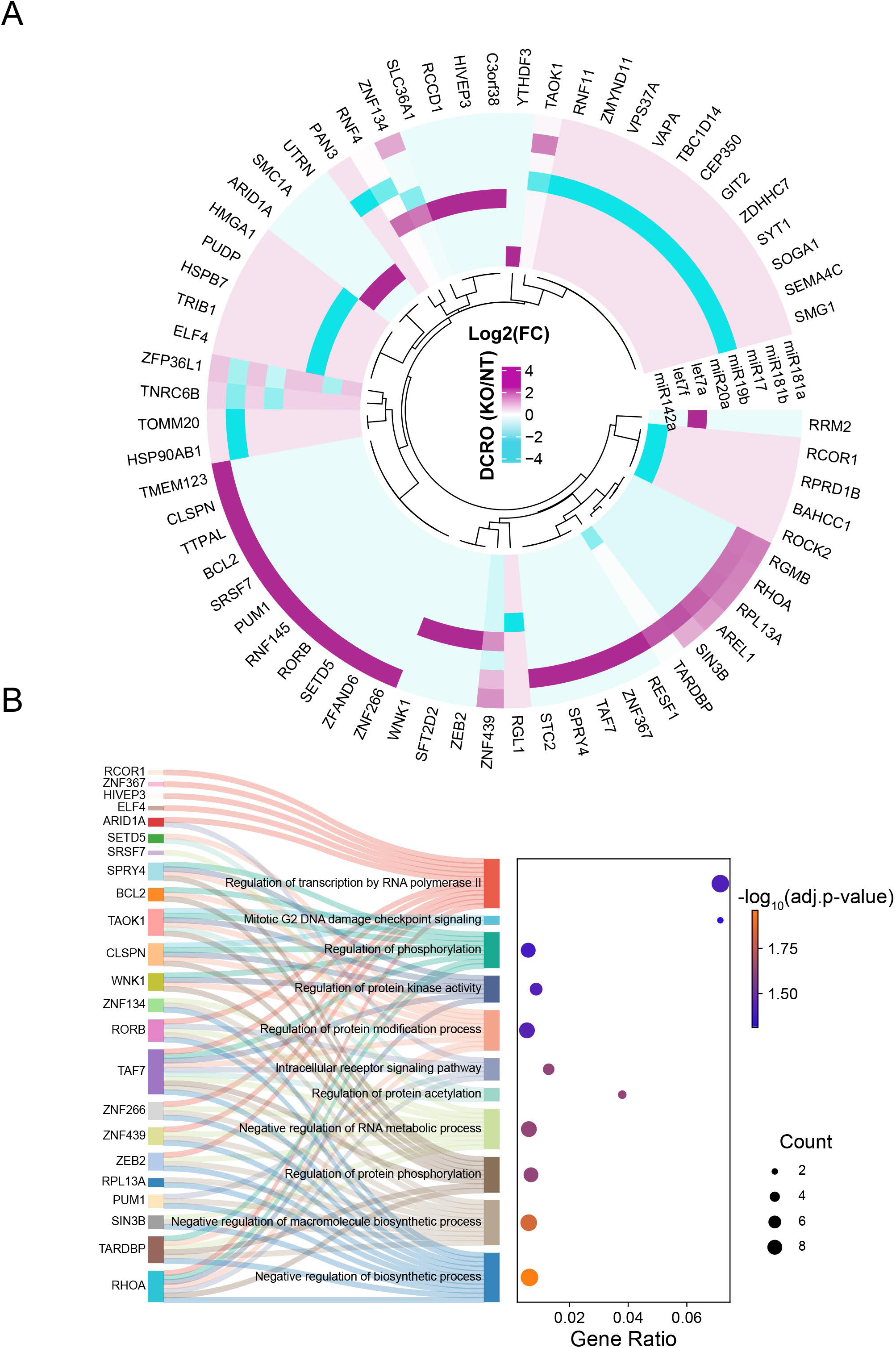
(A) Circle plot depicting Differential Chimeric Read Occupancy (DCRO) of miR-181a/b, miR-17, miR-19b, miR-20, let-7a/f and miR-142 on I3 target mRNAs, between control (NT) and I3 knock-out (I3KO) cells, with respect to I3KO with a log2FC of +1 and -1 and p value of 0.05. (B) Hallmark enrichment analysis of mRNAs with increased Differential Chimeric Read Occupancy (DCRO) from panel E. Significantly enriched pathways had a FDR of less than 0.05. Genes part of the enriched pathways are annotated and connected by a Sankey plot.

**Supplementary figure 5.**
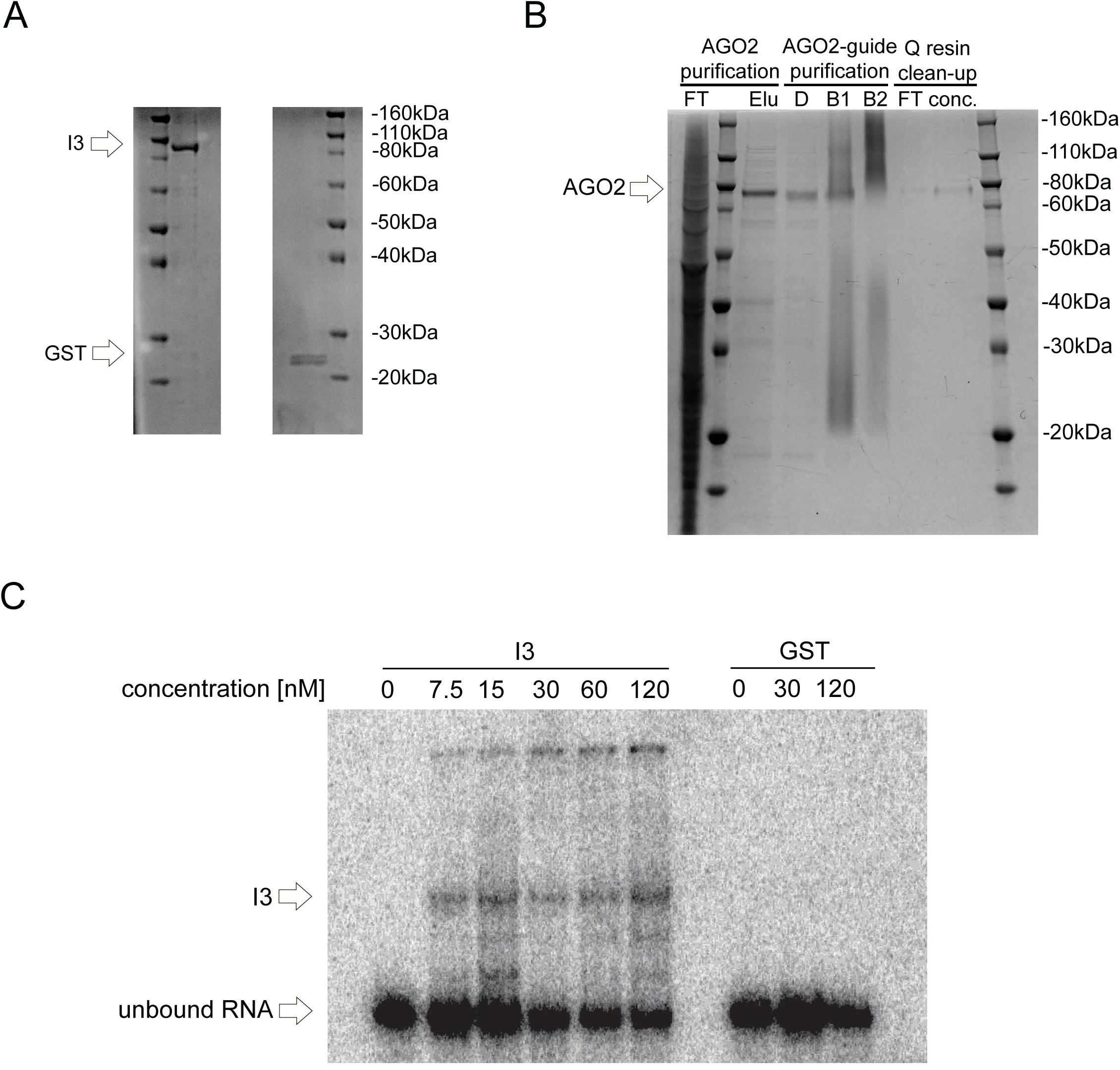
(A) Coomassie-stained gel of recombinantly expressed and purified I3 after FPLC purification to exclude endogenous GST. (B) Coomassie-stained gel of recombinantly expressed and purified AGO2 across the microRNA loading process. FT = flow-through (unbound proteins); Elu = elution (eluted AGO); D = purified AGO after microRNA loading; B1 = bead1 (bound AGO2-let7 to capture resin), B2 = bead2 (capture resin after AGO2-let-7 elution); FT = flow-through during Q-resin clean-up; conc. = concentrated and purified AGO2-let7 complex. (C) Electrophoretic Mobility Shift Assay (EMSA) using recombinant I3 and GST. Partial 3’-UTR sequence from the oncogene *HMGA2* was used as the RNA substrate.

## Notes

### Competing Interest Statement

The authors have declared no competing interest.

